# A global database of insect traits and anthropogenic associations

**DOI:** 10.64898/2026.02.05.704013

**Authors:** E. Manfrini, N. Sauvion, P-O. Maquart, L. Legal, O. Blight, E. Duquesne, C. Hanot, A. Bang, B. Geslin, F.R. Goebel, D. Fournier, Å. Berggren, M. Javal, E. Angulo, S. Pincebourde, M. Zakardjian, J-F. Vayssière, D. Renault, C. Le Lann, S.A.P. Derocles, B. Leroy, F. Courchamp

**Affiliations:** Université Paris Saclay, CNRS, AgroParisTech, Ecologie Société Evolution, Gif-sur-Yvette, France; Laboratoire de Biologie des Organismes et des Écosystèmes Aquatiques-BOREA, Muséum national d’Histoire naturelle (MNHN), SU, CNRS, IRD, UA, F-75005 Paris, France; Plant Health Institute of Montpellier (PHIM), Univ. Montpellier, INRAE, CIRAD, Institut Agro, IRD, Montpellier, France; Institut de Recherche pour le Développement, Université Paris-Saclay, UMR-247 Évolution, Génomes, Comportement et Écologie, 91198, Gif-sur-Yvette, France; Université de Toulouse, Toulouse INP, CNRS, IRD, CRBE, Toulouse, France; IMBE, Avignon Université, Aix Marseille Université, CNRS, IRD, Avignon, France; Université libre de Bruxelles, FRS-FNRS, Evolutionary Biology and Ecology, Brussels, Belgium; Biology Group, School of Arts and Sciences, Azim Premji University, Bhopal, 462002, India; Univ Rennes, CNRS, Ecobio [(Ecosystème, Biodiversité, Evolution)] – UMR, 6553 Rennes, France; Univ Montpellier, CIRAD, Montpellier, France; Department of Ecology, Swedish University of Agricultural Sciences, Uppsala, Sweden; CBGP, IRD, CIRAD, INRAE, Montpellier SupAgro, Univ Montpellier, Montpellier, France; Estación Biológica de Doñana (EBD-CSIC), 41092 Sevilla, Spain; Université de Tours, Parc Grandmont, 37200 Tours, France; IMBE, Aix Marseille Université, CNRS, IRD, Avignon Université, Marseille, France

**Keywords:** AnthopInsect, edible insects, invasive species, ecological traits, macroecological indicators, functional descriptors

## Abstract

Insect research remains hindered by limited data availability and fragmented knowledge compared to other, better-documented taxonomic groups. Yet, both the macroecological and the insect research communities highlight the need to integrate large-scale ecological trait datasets for insects. Insects and humans are interconnected through diverse relationships, ranging from beneficial interactions, such as the use of insects as nutritional resources, to adverse impacts, including their role in to the emergence and spread of biological invasions. Understanding the traits of insects associated with human use, movement and impact is therefore important for linking insects to ecosystem function and global change. We present *AnthropInsect*, one of the largest database on insect traits to date, which uniquely includes variables describing human-insect associations. *AnthropInsect* describes species through 35 variables grouped into five categories: (i) taxonomic descriptors; (ii) ecological descriptors (native bioregions and habitat); (iii) human-insect associations (edibility and invasive status); (iv) functional traits (behavior, morphology, life history and feeding); (v) and macroecological descriptors of native-range geography and climate. *AnthropInsect* currently includes 5867 species across six major orders: Coleoptera, Lepidoptera, Hemiptera, Hymenoptera, Orthoptera and Blattodea. Data extracted from peer-reviewed, grey literature and from existing databases were standardized and validated with expert knowledge to ensure accuracy. By providing traits data with information on insect–human interactions, this rigorously curated resource supports global research in entomology, ecology, conservation, and global change.

## Background & Summary

Over the last decades, there has been increasing recognition and consensus that biodiversity is multi-dimensional, encompassing taxonomic, phylogenetic, and functional diversities, all of which are essential to consider for effective conservation strategies^1–4^. Trait-based approaches have therefore become central in ecology and are now applied across a broad range of taxa^5–8^. Large curated and harmonized trait databases have been developed, enabling comparative studies of macroecological patterns and trends. These initiatives have focused particularly on plants and vertebrates, with notable examples including TRY, a global database of plant traits^9^; AmphiBIO, which compiles natural history traits for amphibians worldwide^10^; FISHMORPH, a database of fish morphological traits^11^; COMBINE, a consolidated database of intrinsic and extrinsic mammal traits^12^; or AVONET, which provides comprehensive functional trait data for all bird species^13^. In contrast, other taxonomic groups, particularly insects, still suffer from major gaps in trait information and understanding of their ecological functions, a limitation known as the Raukiærian shortfall^14–18^.

In addition to being the most speciose and abundant class of animals, insects are also functionally diverse and fulfil key ecological roles, including pollination, nutrient cycles, seed dispersal, and ecosystem engineering^19,20^. Through their diverse trophic functions (e.g., herbivores, decomposers, predators, and parasitoids), functional niches and traits, they regulate plant and arthropod populations and constitute a strong basis of many trophic networks^21–23^. As these ecological functions are essential to ecosystem functioning and human well-being, insects represent a priority group for macroecological and conservation studies^24^. In response, substantial efforts have been made by the scientific community and initiatives have generated order-, family-, region-, or ecosystem-specific insect trait datasets. These include datasets on North American lotic insects traits and their evolutionary associations^25^; ants traits with associated georeferenced assemblage-level data at the global scale^26^; odonates phenotypic traits^27^; European aquatic macroinvertebrates and their dispersal traits^28^; heteroptera traits from a protected area of China^29^; and bees^30^. However, existing resources remain highly specific and insufficiently integrated, limiting cross-taxonomic and cross-regional comparisons, while available trait data remain fragmented and lack centralization^26,27^. To address this challenge, Cardoso et al.^31^ called for the establishment of a global network of collaborators to collectively accelerate the compilation of insect trait data and develop a unified insect trait database, providing guidelines and a framework for such a database.

Insects are often closely linked to and influenced by anthropogenic activities, with interactions that can have both positive and negative effects on human well-being. On the one hand, they can provide valuable resources, acting as biological control agents of pests or contributing to medical applications^32,33^. On the other hand, some insects play key roles in global change processes, which are major drivers of global biodiversity loss. Among these processes, biological invasions - defined here as the human-mediated introduction of species beyond their historical ranges - represent a major driver of global biodiversity loss^4,34,35^. Insects are the most frequently introduced taxonomic group and one of the most invasive taxa globally^36–38^. Their introductions can have substantial impacts on ecosystems, human well-being and health, and economic activities^39,40^. Insect introductions are increasing with globalization, many of which are linked to human use and trade. This tight relationship is particularly important in the case of farmed or commercially exploited species that are often selected for traits including rapid growth, early reproduction, high fecundity, generalist diets, disease resistance, and broad environmental tolerance^41,42^. These traits often predispose insect species to successfully establish and spread in novel regions thus making trait-based studies critical for information on the likely success of invasion and for management strategies^43–45^. This illustrates the tight human-insect associations and the need for an adequate and comprehensive insect trait database.

With the recent rise of insect farming for food and feed, several studies have highlighted the risk that edible species raised for industrial purposes could contribute to new biological invasions when farmed outside their native ranges^46–48^. We created *AnthropInsect*, a database focused on insect species relevant to human activities, illustrated here through both negative (invasive species) and positive (edible species) associations. This database provides trait coverage for 5867 species spanning six of the most speciose insect orders (Coleoptera, Lepidoptera, Hemiptera, Hymenoptera, Orthoptera, and Blattodea)^49^. It compiles 35 variables describing taxonomic, ecological, functional, human-association and macroecological dimensions, which are often linked to the processes driving biological invasions^45,50^. *AnthropInsect* is a standalone database that encompasses key dimensions for understanding insect traits in an increasingly human-dominated world. It is also interoperable and can naturally complement and be integrated with the Global Repository of Insect Traits (GRIT) initiative proposed by Cardoso et al.^31^. As an open, evolving, and community-driven resource, *AnthropInsect* is designed to support large-scale comparative studies in entomology, invasion biology, climate change research, functional ecology, and applied sciences.

## Methods

This section describes how the dataset was assembled, including the rationale for insect species inclusion and the main steps used to compile, curate, and validate records.

### Insect species selection

*AnthropInsect* currently covers two important groups of species associated with human activities: species that are raised and known to be consumed by humans (hereafter referred to as edible species), and species reported to be transported and introduced by humans (hereafter referred to as alien insect species). Species records were extracted from existing databases or from specific sources (recorded in the “Source” column).

We selected edible insect species from the list compiled by Van Itterbeeck & Pelozuelo.^51^, derived from Jongema.^52^, which provides comprehensive information of the worldwide usage of insects. Where possible, we refined usage categories and retained only species listed as “food insect species”. For the first version of the *AnthropInsect* database, we focused on six of the most represented edible insect orders - Coleoptera, Lepidoptera, Hemiptera, Hymenoptera, Orthoptera, and Blattodea - which together account for over 90% of reported edible insects. This yielded 1,276 edible insect species.

For alien insect species, we distinguished two categories. *Established non native species* (hereafter referred to as ‘*Established*’) are defined as introduced taxa that maintain viable, self-sustaining populations outside their native range. *Invasive non native species* (hereafter referred to as ‘*Invasive*’) are established taxa that have spread and cause documented environmental and/or socio-economic impacts^53^. We first compiled established insect species from Seebens et al.^54^, which documents taxa with a known arrival date in at least one non-native region. Within the six target orders (see previous paragraph), we identified 3,123 *Established* insect species, representing approximately 75% of all *Established* insect species reported by Seebens et al.^54^. Second, to assemble a representative list of *Invasive* insects, we combined information from three major sources: the Global Invasive Species Database (GISD, http://www.iucngisd.org/gisd/), the Global Register of Introduced and Invasive Species (GRIIS^55^) and the CABI Invasive Species Compendium (https://www.cabidigitallibrary.org/product/qi). These resources, differing in both spatial coverage and species composition, improved overall database completeness. Within the six target orders, we identified 1,467 *Invasive* insect species, corresponding to approximately 70% of all known *invasive* insect species within the stated databases. In *AnthropInsect*, we labeled this invasive status to species that are “*Invasive*” in at least one region worldwide.

To ensure taxonomic consistency, we validated insect species names from the three lists against the GBIF taxonomic backbone^56^ (using the R package rgbif^57^). When automated matching failed, order-specific taxonomic experts performed manual validation. We removed duplicate entries to retain unique species records in the final dataset.

Finally, we merged insect species from the lists into a unified checklist. We reclassified species initially ‘*Established’* as ‘*Invasive’* when they were documented as invasive in at least one of the invasive-species sources. We classified insect species that were absent from both alien categories (*i*.*e*., not reported as established or invasive outside their native bioregions in the source databases) as “*Native only*”. As of today, the final dataset includes 5867 insect species (**Table 2**).

### Selection of variables and trait data extraction

We described each insect species in the database using 35 variables organised into five categories (**Table 1**): (i) taxonomic descriptors (9 variables; order to species level, including the GBIF backbone taxon ID); (ii) ecological descriptors (2 variables; native bioregions, and larval and adult habitats); (iii) human associations (2 variables; edibility and invasion status); (iv) functional traits (18 variables) describing life histories, behavior, feeding, and morphology; and (v) macroecological variables (4 variables) capturing native-range geographic distribution and climatic conditions (temperature and humidity).

**Table 1:** Definitions and categories of insect species variables in the *AnthropInsect* database.

| Variables | Definition | Categories |
| --- | --- | --- |
| Identification |  |  |
| scientificNameID | GBIF backbone taxon ID. |  |
| Source | Database or study names where invasive status or species edibility were found. |  |
| Taxonomy |  |  |
| Order<br>Suborder<br>Infraorder<br>Family<br>Superfamily<br>Genus<br>Scientific name<br>Species | Species taxonomy. |  |
| Ecological descriptors |  |  |
| Native bioregion | Native species territory whose limits are defined by geographical limits that take into account both human communities and ecosystems (based on vertebrate bioregions from Holt et al.<sup>62</sup> , as insect bioregions are not yet delineated in the literature). | Nearctic; Neotropical; Palearctic; Afrotropical; Oriental; Australian ; Oceanian; Antarctic; Marine; Panamanian; Sino-Japanese; Saharo-Arabian; Antarctica |
| Habitat larva | Habitat where the life stage cycle occurred. | Aquatic; Semi-aquatic; Terrestrial |
| Habitat adult |  | Aquatic; Semi-aquatic; Terrestrial |
| Human associations |  |  |
| Edibility | Species known to be consumed by humans (adapted from Van Itterbeeck & Pelozuelo<sup>51</sup>). | YES; NO |
| Invasive status | Native only species: species that occur naturally within their original geographic range and have not been transported by human activities. | Native only; Established; Invasive |
|  | Established species: species that humans have introduced outside their native range and that have successfully formed viable, self-sustaining populations, potentially spreading beyond the initial introduction site.<br>Invasive species: established species that spread widely and cause negative impacts on native biodiversity and socio economic systems. |  |
| **Functional traits** |  |  |
| ***Life history*** |  |  |
| **Metamorphosis** | The changes in form that occur during the insect cycle. | Holometabolous ; Hemimetabolous |
| **Inactivity period** | Temporary state of inactivity that is typically induced by external factors such as unfavorable environmental conditions (include diapause and quiescence). | YES; NO |
| **Host** | Strict relationship between the insect species and another species for its life cycle, reproduction, or survival. | YES; NO |
| **Host specificity** | Scientific name of the taxonomic level of the host. |  |
| **Parthenogenesis** | Can produce offspring through asexual reproduction. | YES; NO |
| **Egg laying pattern** | Details the egg-laying patterns: all at once, continuously (along adult's life), one by one or in clusters (patches). | Once; Continuously; Singly; Clusters |
| **Parental care** | Behavior exhibited by adult insects to care for their offspring (e.g., eggs and larvae), involving the adults' presence before and/or after the eggs hatch to enhance the offspring's survival and growth. | YES; NO |
| **Lifespan** | Duration of an individual insect's life, measured from the egg stage to death. For insects living in colonies, consider the lifespan of the queen. | Quantitative data in days |
| **Voltinism** | Number of generations an organism completes within a single year. | Quantitative data without unit |
| **Fecundity** | Number of eggs laid by a female over its lifespan. For insects living in colonies and showing reproductive skew, consider the number of eggs laid by the major reproductive or the queen. |  |
| **Dispersal mode** | Species locomotion for moving or spreading. For insects living in colonies, consider the locomotion of the queen. | Flying; Swimming; Walking; Burrowing; Skating; Abiotic dispersal |
| ***Behavior*** |  |  |
| **Sociability** | The degree to which individuals within a population engage in social behaviors or form social structures. | Solitary; Social (includes communal, quasisocial, subsocial); Eusocial |
| **Aggressiveness** | The behavior or tendency of individuals to display aggression, which involves actions intended to harm or intimidate others. | NO; Intra-specific; Inter-specific; Intra-specific/Inter-specific |
| ***Feeding*** |  |  |
| **Diet larva** | Diet of the life stage of the insect species. | Phytophage; Mycophage; Saprophage; Zoophage; Omnivore |
| **Diet adult** |  | Phytophage; Mycophage; Saprophage; Zoophage; Omnivore; Nothing |
| **Feeding behavior larva** | Location of the feeding area during the larval or adult stage. | Endophage; Exophage |
| **Feeding behavior adult** |  | Endophage; Exophage; Nothing |
| ***Morphology*** |  |  |
| **Body size adult** | Adult body length. For insects living in colonies, consider the queen's size. | Quantitative data in millimeter |
| **Macroecological variables** |  |  |
| ***Species known occurrences*** |  |  |
| **Occurrence numbers** | Number of species occurrences from 1980 to 2024. | Quantitative data |
| ***Distribution ranges*** |  |  |
| **Latitude range** | Latitudinal species distribution in its native range. | Quantitative data in decimal degree |
| **Longitude range** | Longitudinal species distribution in its native range. |  |
| ***Climatic condition ranges*** |  |  |
| **Thermal range** | Thermal range (bio5 - bio 6) of a species in its native range.<br>Bio5 : Mean daily maximum air temperature of the hottest month.<br>Bio6 : Mean daily minimum air temperature of the coldest month. | Quantitative data in Celsius degree |
| **Humidity range** | Humidity range of a species in its native range.<br>Hurs_max : Maximum monthly near surface relative humidity.<br>Hurs_min : Minimum monthly near surface relative humidity. | Quantitative data in % of humidity |

**Table 2:** Categorization of insect species by edibility and invasive status, the two criteria motivating the development of *AnthropInsect*.

|  |  | Edibility |  |  |
| --- | --- | --- | --- | --- |
|  |  | YES | NO | Total |
| Invasive status | Native only | 1185 | 92 | 1277 |
|  | Established | 30 | 3093 | 3123 |
|  | Invasive | 61 | 1406 | 1467 |
| Total |  | 1276 | 4560 | 5867 |

With the exception of macroecological variables, we compiled trait information primarily from peer-reviewed and grey literature sources. While this approach demanded substantial taxonomic expertise, it greatly contributed to improving data completeness and consistency across the database, with various experts also contributing their own unpublished field data.

We assigned traits values using three documented approaches: (i) direct extraction: we extracted values from the literature and, where available, from existing databases (e.g.,^58–60^); (ii) expert assignment: we assigned values based on expert knowledge for variables that are well documented and conservative at higher taxonomic levels (e.g., metamorphosis type at the order or family level); and (iii) phylogenetic inference: we inferred values from the closest related taxa when species-level information was unavailable, prioritising congeners (same genus). We applied this inference mainly to Orthoptera and only rarely to other groups. For each record, the database reports the assignment method and the corresponding source information to ensure full traceability.

We compiled macroecological variables using an automated two-step workflow. First, we downloaded occurrence records from GBIF for the period 1980-2024 using the R package *rgbif* ^57^. Second, we extracted temperature and humidity values from Climatologies at High Resolution for the Earth’s Land Surface Area (CHELSA, 1 km pixel resolution; chelsa-climate.org^61^) at each occurrence location. To ensure consistent comparisons among species with multiple invasion-status categories, we restricted native-range macroecological descriptors to occurrence records within each species’ native bioregions. Additionally, we excluded records located outside mainlands (i.e., in marine environments). Lastly, we computed four native-range macroecological descriptors (**Table 1**): (i) latitude range, defined as the difference between maximum and minimum latitudes (decimal degrees); (ii) longitude range, defined as the difference between maximum and minimum longitudes (decimal degrees); (iii) thermal range, defined as the difference between maximum and minimum temperatures (°C; bio5 and bio6) extracted from occurrence locations corresponding to the hottest and coldest points within the range, respectively; and (iv) humidity range, defined as the difference between maximum and minimum humidities (%, hurs_max and hurs_min) extracted from occurrence locations corresponding to the wettest and driest points of the range, respectively. For species with more than 10 retained occurrences, we reduced the influence of extreme values by trimming the lowest 1% and highest 1% of climatic values prior to summary calculations.

### Database completeness

Overall, 51% of all species in the database have complete data, with *Native only* species showing the highest completeness (78%), followed by *Invasive* species (51%) and *Established* species (41%) (**Figure 1**). This pattern partly reflects our current prioritisation of data compilation efforts toward edible insect species, most of which fall into this *Native only* category. *Sociability, Aggressiveness, Metamorphosis*, and *Parthenogenesis* are among the traits documented for the largest number of species in *AnthropInsect* (**Figure 1**). These variables are often conserved within closely related taxa, which allowed values to be assigned with confidence at higher taxonomic levels (e.g., family or order) when species-level information was unavailable. By contrast, quantitative traits that required manual entry are the least complete (*lifespan, fecundity, voltinism*, and *body size adult*), as they are typically highly species-specific and are not consistently reported across the literature. Macroecological variables derived from occurrence data (native *latitude* and *longitude ranges*, and native *thermal* and *humidity ranges)* could be computed for less than 50% of species. This primarily reflects limited availability of occurrence records in GBIF, and, for established and invasive species, insufficient information on native bioregions, which prevents identification of the native range within occurrence data and derivation of associated distribution and climatic metrics^14,16^. Despite incomplete coverage of native bioregions, *AnthropInsect* includes species originating from all major global regions (**Figure 2**).

**Figure 1:**
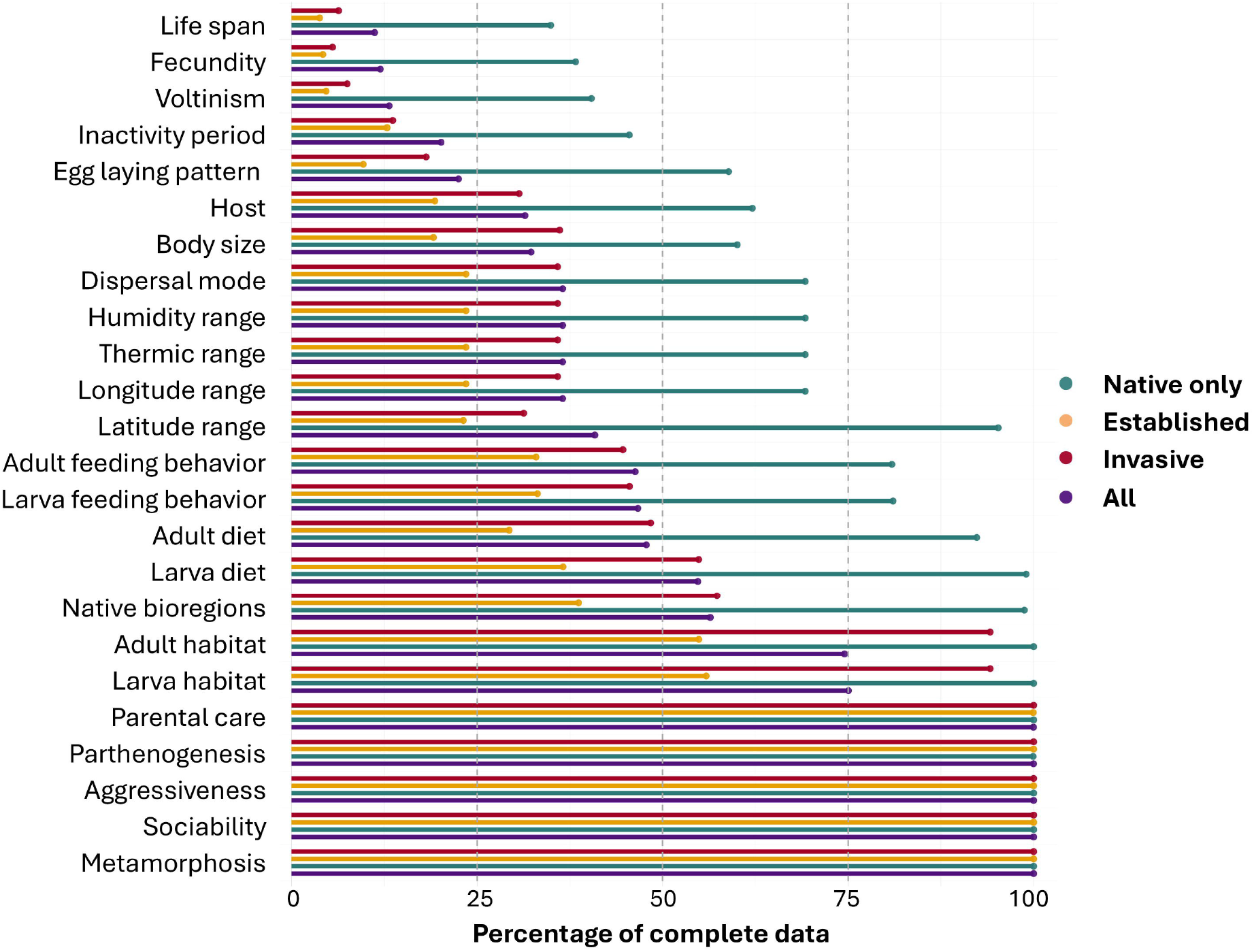
Percentage of complete data per variable and per invasive insect species status. Violet bars represent all species, green bars *Native only* species, yellow bars *Established* species, and red bars *Invasive* species. See **Table 1** for the definition of the variables.

**Figure 2:**
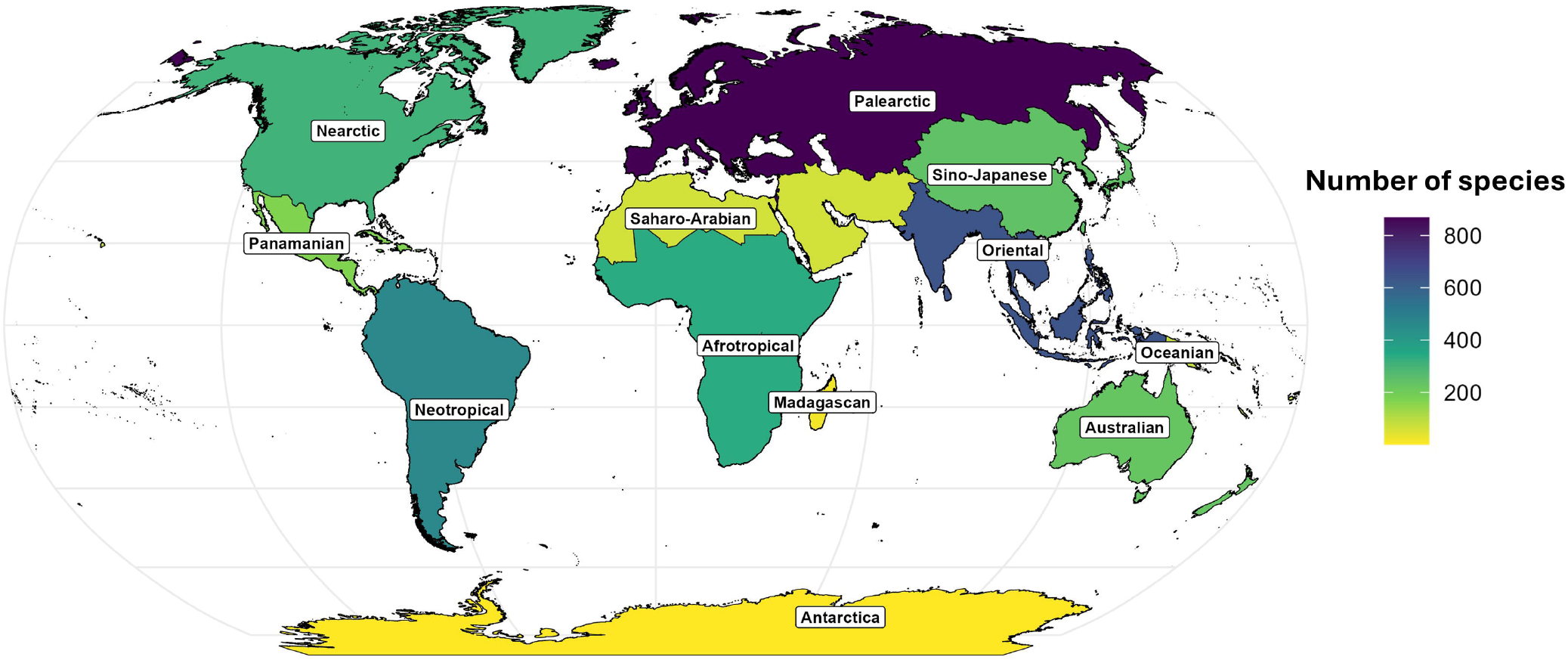
Number of species from *AnthropInsect* across inferred native bioregions^62^.

### Possible application

By integrating insect trait data with human-related and macroecological information, *AnthropInsect* fills critical knowledge gaps across a wide range of disciplines, including entomology, invasion biology, climate change, functional ecology, and applied sciences. *AnthropInsect* follows the FAIR principles which make it readily integrable into the larger Global Repository of Insect Traits initiated by Cardoso et al.^31^, thereby contributing to a global collaborative network aimed at accelerating the compilation of insect trait data and improving transparency, accessibility, and data sharing within the field.

Currently, the database focuses on edibility and invasive status, which are likely among the most important species groups. However, human associations span a broader scope of possible species groups, which can be integrated in the database in the future. These include medicinal uses, agricultural (native) pests, bioindicators, disease vectors, cultural significance (e.g., ecotourism, symbolism), ecosystem services, and material uses (e.g., honey, silk, dyes, wax).

## Data records

The current release of *AnthropInsect* comprises 5867 insect species described by 35 variables grouped into five categories: (i) taxonomic descriptors, (ii) ecological descriptors, (iii) human associations, (iv) functional traits, and (v) macroecological variables.

The *AnthropInsect* database is publicly available to facilitate access and reuse. It has been deposited in Data InDoRES^49^ (https://doi.org/10.48579/PRO/LYB70N) as a public, community-recognized repository. It is a spreadsheet organised into five tabs that document the database structure, metadata, and traceability. The first tab provides an overview of the database (e.g., number of species, variables, etc.) and describes the content of each tab in the file (labelled ‘Read.me’). The second lists experts and contributors which are involved in data compilation, together with their taxonomic expertise (labelled ‘Authors_Tax-Repartition’). The third tab (‘Variables’) provides variable names, definitions, and categories. The *AnthropInsect* dataset is stored in the ‘Database’ tab, while an additional tab (‘ References’) contains the complete list of reference IDs corresponding to the sources used for trait extraction. Because *AnthropInsect* can be expanded collaboratively, we provide a ‘Completion Guidelines’ document that describes recommended procedures and quality-control steps to ensure consistent future database updates.

All data associated with this project, including the *AnthropInsect* spreadsheet are publicly available on Data InDoRES^49^. Scripts used to (1) extract macroecological variables, (2) automate the completion of selected database fields, and (3) support data visualisation are available on GitHub (https://github.com/Elena-Manfrini/AnthropInsect).

### Technical Validation

We implemented a series of checks to ensure the reliability and interoperability of *AnthropInsects*. Trait definitions and coding followed published recommendations for standardised trait reporting and harmonisation (e.g.,^31,63^). To maintain up-to-date taxonomic classifications and facilitate interoperability with external biodiversity resources, species names were validated against the GBIF taxonomic backbone, and we recorded the corresponding GBIF identifiers (e.g., accepted name usage IDs where available^31^). We also cross-referenced and integrated information from existing trait databases (e.g.,^58–60^) to improve completeness and consistency.

To support traceability, we associate each trait value with a source and provide a complete list of reference identifiers used for data extraction. Contributor and expert information is documented to facilitate follow-up in case of questions or potential errors. Finally, all automated data retrieval and completion steps (e.g., occurrence downloads, climate extraction, and scripted field completion) are documented and implemented through version-controlled R scripts in the repository, ensuring that key processing steps are transparent and reproducible (https://github.com/Elena-Manfrini/AnthropInsect).

### Usage notes

*AnthropInsect* is intended to support a broad range of users in entomology and ecology, particularly for studies on biological invasions, functional diversity, and human-insect associations. The complete dataset is publicly available to facilitate access and reuse^49^.

We encourage corrections, updates, and new contributions through a standardised workflow. Contributions can be submitted via our email address and should provide new and/or revised information in a spreadsheet format consistent with the database structure (e.g., identical columns and controlled vocabularies), as well as any new columns related to human–insect associations. Detailed information on the *AnthropInsect* database, along with instructions for data completion and common mistakes to avoid, are provided in the ‘Completion Guidelines’ document^49^.

## Code availability

The R scripts used for extracting macroecological data and to visualize data from the *AnthropInsect* database are available in the GitHub repository (https://github.com/Elena-Manfrini/AnthropInsect).

## Data availability

The *AnthropInsect* database is deposited in Data InDoRES^49^ (https://doi.org/10.48579/PRO/LYB70N).

## Acknowledgements

This study received support from the AXA Research Fund Chair for Biological Invasions at the University of Paris Saclay.

## Author contributions

E.M, F.C, B.L conceived the initial project and designed the study strategy. E.M. managed the extensive teamwork. E.M designed the database. All the authors contributed to improving the database quality, performed the literature searches, material screening and data collation. E.M. developed the scripts for analyses. E.M. and A.B. took the lead in writing the paper with inputs and reviews from all contributing authors. All authors read and approved the final manuscript.

## Bibliography

1. Strecker, A. L., Olden, J. D., Whittier, J. B. & Paukert, C. P. Defining conservation priorities for freshwater fishes according to taxonomic, functional, and phylogenetic diversity. Ecological Applications 21, 3002–3013 (2011).

2. Mazel, F. et al. Multifaceted diversity–area relationships reveal global hotspots of mammalian species, trait and lineage diversity. Global Ecology and Biogeography 23, 836–847 (2014).

3. Brum, F. T. et al. Global priorities for conservation across multiple dimensions of mammalian diversity. Proc. Natl. Acad. Sci. U.S.A. 114, 7641–7646 (2017).

4. IPBES, Brondizio, E., Diaz, S., Settele, J. & Ngo, H. T. Global Assessment Report on Biodiversity and Ecosystem Services of the Intergovernmental Science-Policy Platform on Biodiversity and Ecosystem Services. 1144 https://zenodo.org/records/6417333 (2019) doi:10.5281/ZENODO.3831673.

5. Chapple, D. G., Simmonds, S. M. & Wong, B. B. M. Can behavioral and personality traits influence the success of unintentional species introductions? Trends in Ecology & Evolution 27, 57–64 (2012).

6. Green, J. L., Bohannan, B. J. M. & Whitaker, R. J. Microbial Biogeography: From Taxonomy to Traits. Science 320, 1039–1043 (2008).

7. Pey, B. et al. Current use of and future needs for soil invertebrate functional traits in community ecology. Basic and Applied Ecology 15, 194–206 (2014).

8. Shipley, B. et al. Reinforcing loose foundation stones in trait-based plant ecology. Oecologia 180, 923–931 (2016).

9. Kattge, J. et al. TRY plant trait database – enhanced coverage and open access. Global Change Biology 26, 119–188 (2020).

10. Oliveira, B. F., São-Pedro, V. A., Santos-Barrera, G., Penone, C. & Costa, G. C. AmphiBIO, a global database for amphibian ecological traits. Sci Data 4, 170123 (2017).

11. Brosse, S. et al. FISHMORPH: A global database on morphological traits of freshwater fishes. Global Ecol. Biogeogr. 30, 2330–2336 (2021).

12. Soria, C. D., Pacifici, M., Di Marco, M., Stephen, S. M. & Rondinini, C. COMBINE: a coalesced mammal database of intrinsic and extrinsic traits. Ecology 102, e03344 (2021).

13. Tobias, J. A. et al. AVONET: morphological, ecological and geographical data for all birds. Ecology Letters 25, 581–597 (2022).

14. Hortal, J. et al. Seven Shortfalls that Beset Large-Scale Knowledge of Biodiversity. Annu. Rev. Ecol. Evol. Syst. 46, 523–549 (2015).

15. Marshall, L. et al. Understanding and addressing shortfalls in European wild bee data. Biological Conservation 290, 110455 (2024).

16. Cardoso, P., Erwin, T. L., Borges, P. A. V. & New, T. R. The seven impediments in invertebrate conservation and how to overcome them. Biological Conservation 144, 2647–2655 (2011).

17. Stork, N. E. How Many Species of Insects and Other Terrestrial Arthropods Are There on Earth? Annu. Rev. Entomol. 63, 31–45 (2018).

18. Troudet, J., Grandcolas, P., Blin, A., Vignes-Lebbe, R. & Legendre, F. Taxonomic bias in biodiversity data and societal preferences. Sci Rep 7, 9132 (2017).

19. Noriega, J. A. et al. Research trends in ecosystem services provided by insects. Basic and Applied Ecology 26, 8–23 (2018).

20. Dangles, O. & Casas, J. Ecosystem services provided by insects for achieving sustainable development goals. Ecosystem Services 35, 109–115 (2019).

21. Yang, L. H. & Gratton, C. Insects as drivers of ecosystem processes. Current Opinion in Insect Science 2, 26–32 (2014).

22. Verma, R. C. et al. The Role of Insects in Ecosystems, an in-depth Review of Entomological Research. IJECC 13, 4340–4348 (2023).

23. Schoenly, K., et al. On the Trophic Relations of Insects: A Food-Web Approach. The American Naturalist 137, 597–638 (1991).

24. Harvey, J. A. et al. Scientists’ warning on climate change and insects. Ecological Monographs 93, e1553 (2023).

25. Poff, N. L. et al. Functional trait niches of North American lotic insects: traits-based ecological applications in light of phylogenetic relationships. Journal of the North American Benthological Society 25, 730–755 (2006).

26. Parr, C. L. et al. GlobalAnts : a new database on the geography of ant traits (Hymenoptera: Formicidae). Insect Conserv Diversity 10, 5–20 (2017).

27. Waller, J. T., et al. The odonate phenotypic database, a new open data resource for comparative studies of an old insect order. Sci Data 6, 316 (2019).

28. Sarremejane, R. et al. DISPERSE, a trait database to assess the dispersal potential of European aquatic macroinvertebrates. Sci Data 7, 386 (2020).

29. Gao, S. et al. A morphological traits dataset of Heteroptera sampled in biodiversity priority areas of Southwest China. Sci Data 11, 694 (2024).

30. Aubouin, L. et al. BeeFunc, a comprehensive trait database for French bees. Sci Data 12, 1302 (2025).

31. Cardoso, P. et al. Toward a global repository of insect traits ( GRIT). Insect Conserv Diversity icad.70035 (2025) doi:10.1111/icad.70035.

32. Pervez, A. & Omkar. Ecology and biological control application of multicoloured Asian ladybird, Harmonia axyridis: A review. Biocontrol Science and Technology 16, 111–128 (2006).

33. Costa-Neto, E. M. Entomotherapy, or the Medicinal Use of Insects. Journal of Ethnobiology vol. 25 (2005).

34. Mathakutha, R. et al. Invasive species differ in key functional traits from native and non‐invasive alien plant species. J Vegetation Science 30, 994–1006 (2019).

35. Zhao, Z. et al. The world’s 100 worst invasive alien insect species differ in their characteristics from related non‐invasive species. Journal of Applied Ecology 60, 1929–1938 (2023).

36. Diagne, C. et al. High and rising economic costs of biological invasions worldwide. Nature 592, 571–576 (2021).

37. Seebens, H. et al. Global rise in emerging alien species results from increased accessibility of new source pools. Proc. Natl. Acad. Sci. U.S.A. 115, (2018).

38. Bertelsmeier, C. et al. Temporal dynamics and global flows of insect invasions in an era of globalization. Nat. Rev. Biodivers. 1, 90–103 (2025).

39. Renault, D. et al. The magnitude, diversity, and distribution of the economic costs of invasive terrestrial invertebrates worldwide. Science of The Total Environment 835, 155391 (2022).

40. Kenis, M. et al. Ecological effects of invasive alien insects. Biol Invasions 11, 21–45 (2009).

41. Van Huis, A. Potential of Insects as Food and Feed in Assuring Food Security. Annu. Rev. Entomol. 58, 563–583 (2013).

42. Ricciardi, A. et al. Invasion Science: A Horizon Scan of Emerging Challenges and Opportunities. Trends in Ecology & Evolution 32, 464–474 (2017).

43. Mally, R. et al. Insect invasion success depends on taxon and trophic group. NB 99, 19–43 (2025).

44. Novoa, A. et al. Invasion syndromes: a systematic approach for predicting biological invasions and facilitating effective management. Biol Invasions 22, 1801–1820 (2020).

45. Renault, D., et al. Environmental Adaptations, Ecological Filtering, and Dispersal Central to Insect Invasions. Annu. Rev. Entomol. 63, 345–368 (2018).

46. Berggren, Å., et al. Approaching Ecological Sustainability in the Emerging Insects-as-Food Industry. Trends in Ecology & Evolution 34, 132–138 (2019).

47. Bang, A. & Courchamp, F. Industrial rearing of edible insects could be a major source of new biological invasions. Ecology Letters 24, 393–397 (2021).

48. Manfrini, E., et al. Preventing the next invasion: Lessons from aquaculture for the safe expansion of insect farming. Journal of Applied Ecology 63, e70311 (2026).

49. Manfrini, E., Sauvion, N., Maquart, P-O., Legal, L., Blight, O., Duquesne, E., Hanot, C., Bang, A., Geslin, B., Goebel, F-R., Fournier, D., Berggren Å., Javal, M., Angulo, E., Pincebourde, S., Zakardjian, M., Vayssière, J-F., Renault, D., Le Lann, C., Derocles, S.A.P., Leroy, B., Courchamp, F. A global database of insect traits and anthropogenic associations. 10.48579/PRO/LYB70N (2026). A global database of insect traits and anthropogenic associations. data.InDoRES (2026) https://doi.org/10.48579/PRO/LYB70N.

50. Cuthbert, R. N. et al. Harnessing traits to predict economic impacts from biological invasions. Trends in Ecology & Evolution S0169534725000886 (2025) doi:10.1016/j.tree.2025.03.016.

51. Itterbeeck, J. V. & Pelozuelo, L. How Many Edible Insect Species Are There? A Not So Simple Question. Diversity, 14(2), 143.

52. Jongema, Y. List of edible insects of the world. (2017).

53. IPBES. Thematic Assessment Report on Invasive Alien Species and Their Control of the Intergovernmental Science-Policy Platform on Biodiversity and Ecosystem Services. Roy, H. E., Pauchard, A., Stoett, P., and Renard Truong, T. (Eds.). (2023) 10.5281/zenodo.7430682.

54. Seebens, H. et al. No saturation in the accumulation of alien species worldwide. Nat Commun 8, 14435 (2017).

55. Pagad, S., et al. Introducing the Global Register of Introduced and Invasive Species. Sci Data 5, 170202 (2018).

56. GBIF Secreteriat. GBIF Backbone Taxonomy. Checklist dataset. (2023).

57. Chamberlain, S. et al. rgbif: Interface to the Global Biodiversity Information Facility. (2025).

58. Middleton-Welling, J. et al. A new comprehensive trait database of European and Maghreb butterflies, Papilionoidea. Sci Data 7, 351 (2020).

59. Shirey, V. et al. LepTraits 1.0 A globally comprehensive dataset of butterfly traits. Sci Data 9, 382 (2022).

60. Gómez-Suárez, et al. global dataset of native and alien distributions of alien species. Sci Data 12, 1914 (2025).

61. Karger, D. N. et al. Climatologies at high resolution for the earth’s land surface areas. Sci Data 4, 170122 (2017).

62. Holt, B. G. et al. An Update of Wallace’s Zoogeographic Regions of the World. Science 339, 74–78 (2013).

63. Moretti, M. et al. Handbook of protocols for standardized measurement of terrestrial invertebrate functional traits. Functional Ecology 31, 558–567 (2017).

